# An assessment of Nano-RECall: Interpretation of Oxford Nanopore sequence data for HIV-1 drug resistance testing

**DOI:** 10.1101/2022.07.22.501121

**Authors:** Kayla Eileen Delaney, Trevor Ngobeni, Conan K. Woods, Carli Gordijn, Mathilda Claassen, Urvi Parikh, P. Richard Harrigan, Gert Uves van Zyl

## Abstract

**Introduction:** Oxford Nanopore Technologies (ONT) offer sequencing with low-capital-layout sequencing options, which could assist in expanding HIV drug resistance testing to resource limited settings. However, sequence analysis remains time time-consuming and reliant on skilled personnel. Moreover, current ONT bioinformatic pipelines provide a single consensus sequence that is not equivalent to Sanger sequencing, as drug resistance is often detected in mixed populations. We have therefore investigated an integrated bioinformatic pipeline, Nano-RECall, for seamless drug resistance of low read coverage ONT sequence data from affordable Flongle or MinION flow cells.

**Methods:** We compared Sanger sequencing to ONT sequencing of the same HIV-1 subtype C polymerase chain reaction (PCR) amplicons, respectively using RECall and the novel Nano-RECall bioinformatics pipelines. Amplicons were from separate assays a) Applied Biosystems HIV-1 Genotyping Kit (ThermoFisher) spanning *protease (PR)* to *reverse transcriptase (RT) (PR-RT)* (n=46) and b) homebrew *integrase (IN)* (n=21). We investigated optimal read-depth by assessing the coefficient of variation (CV) of nucleotide proportions for various read-depths; and between replicates of 400 reads. The agreement between Sanger sequences and ONT sequences were assessed at nucleotide level, and at codon level for Stanford HIV drug resistance database mutations.

**Results:** The coefficient of variation of ONT minority variants plateaued after a read depth of 400-fold implying limited benefit of additional depth and replicates of 400 reads showed a CV of ∼6 % for a representative position. The average sequence similarity between ONT and Sanger sequences was 99.3% (95% CI: 99.1-99.4%) for PR-RT and 99.6% (95% CI: 99.4-99.7%) for INT. Drug resistance mutations did not differ for 21 *IN* sequences; 16 mutations were detected by both ONT- and Sanger sequencing. For the 46 *PR* and *RT* sequences, 245 mutations were detected by either ONT or Sanger, of these 238 (97.1%) were detected by both.

**Conclusions:** The Nano-RECall pipeline, freely available as a downloadable application on a Windows computer, provides Sanger-equivalent HIV drug resistance interpretation. This novel pipeline combined with a simple workflow and multiplexing samples on ONT flow-cells would contribute to making HIV drug resistance sequencing feasible for resource limited settings.

## Introduction

There is an estimated 37.7 million people living with HIV today. UNAIDS has set ambitious targets to diagnose at least 90%, treat 90% of these and achieve viral load suppression in 90% of those receiving treatment by 2020 and to raise these targets to 95% by 2030[1]. Although the 2020 targets have not been met universally, they emphasize the importance of achieving durable suppression in a large majority of patients. With the increased use of antiretroviral agents for pre-exposure prophylaxis (PrEP) and particularly the use of the same drugs or drug classes in treatment and prophylaxis such as nucleos(t)ide reverse transcriptase inhibitors (NRTIs) especially tenofovir and emtricitabine, non-nucleoside reverse transcriptase inhibitors (NNRTIs) and integrase strand transfer inhibitors (INSTIs), surveillance for increased drug resistance would remain crucial. Sustained suppression is threatened by transmitted or acquired drug resistance. Drug resistance testing is useful in surveillance, as increased transmitted drug resistance may either require HIV drug resistance testing (HIV-DRT) prior to commencing antiretroviral therapy (ART) or a change in first-line treatment recommendations [2]. HIV-DRT is also useful for individual management, especially in informing the choice of regimens in treatment experienced patients with complicated treatment histories[3].

Genotypic HIV-DRT has historically been performed by Sanger Sequencing. It is however a complicated process, with many steps and which require a high level of skill. RECall software has helped to overcome some of these hurdles by providing a freely accessible hands-free interpretation of sequence data[4]. This ensured a lower skills requirement and has ensured uniformity in quality assurance of Sanger Sequencing-based assays for HIV drug resistance in *polymerase (pol)* [4] and of *envelope (env)* for genotypic prediction of tropism[5]. Next generation or third generation sequencing is projected to replace Sanger Sequencing in the near-future, as it has the benefit of detecting low abundance variants (LAV), and could be more cost-efficient than Sanger sequencing when barcoding and pooling multiple samples[6]. However, the high start-up costs, complicated workflow, which require multiple steps, and the long run-time has limited its utility. Oxford Nanopore Technology (ONT) is a long-read third generation sequencing platform and offers several advantages. It offers low start-up cost solutions such as the portable MinION sequencing device with standard flow cells or a cheaper alternative using the Flongle adapter with single-use Flongle flow cells[7]. Moreover, sequence depth could be tailored to one’s needs, by using a “read-until” approach. The major pitfall has been low read accuracy, especially in homopolymer regions, particularly at drug resistance-associated mutations such as K103N and K65R. Moreover, for drug resistance testing, simply obtaining a majority consensus sequence is not enough as drug resistance mutations often occur as mixtures. When dealing with highly conserved genomes such as SARS-CoV-2, the historic low ONT read-accuracy was not prohibitive at the consensus level, when having sufficiently high sequence depth. However, when dealing with highly diverse genomes such as HIV, obtaining accurate alignments have been challenging. Recently we showed an overall agreement of 99.4% between ONT and Illumina when combining a relative low accuracy basecaller with novel iterative alignment pipeline for sequencing HIV proviral PCR products at limiting dilution, with each PCR product expected to represent a single original template[8]. However, in this instance the aim was to obtain a single consensus derived from a single template; such single genome sequencing is however costly and cumbersome. Moreover, available ONT bioinformatics pipelines assume a single consensus, whereas drug resistance mutations often occur as mixed populations. Recent improvements in base-calling combined with the use of a bioinformatics pipeline, specifically developed to correct ONT errors, would make ONT suitable for HIV bulk sequencing. We therefore aimed to assess the agreement of bulk ONT sequencing with Sanger sequencing using a newly developed pipeline, Nano-RECall, while optimizing sequence depth for limited resources. Nano-RECall provides an easy-to-install pipeline on a Windows PC, that does not require bioinformatics skills and which is ideal for settings with limited electronic and/or internet bandwidth resources.

## Methods

The study was a laboratory-based comparison of ONT sequencing and Nano-RECall versus Sanger sequencing. All samples were anonymized routine-diagnostic laboratory samples. A waiver of informed consent was therefore granted. The study had been approved by the Stellenbosch Health Research Ethics Committee N19/07/097.

### Sample material included

Residual samples from routine genotypic HIV drug resistance testing, performed in a public service regional laboratory, in Cape Town, South Africa were included (data collection period: September 2021 to January 2022). Drug resistance testing spanning *protease (PR)* and *reverse transcriptase (RT)* had been performed using Sanger Sequencing with the Applied Biosystems HIV-1 Genotyping Kit (ThermoFisher Scientific). We obtained the most recently generated residual nested-PCR amplicons tested and since amplicon volumes were limited, selected 54 with the highest DNA yield after PCR purification with AMPure XP paramagnetic beads from a total of 65. Similarly, we obtained 35 of the most recent nested-PCR amplicons that had *integrase (IN)* sequencing, using a published in-house method [8] of which 24 with the best DNA yield after PCR purification with the QIAquick PCR purification kit, were included.

### Library preparation of ONT sequencing

Amplicons were included in multiplex runs of 24, each with a different barcode for demultiplexing. The DNA library was prepared by following the instructions from the native barcoding amplicons (with EXP-NBD104, EXP-NBD114, and SQK-LSK109) protocol provided by ONT. Libraries were sequenced with Flongle flow cells and therefore, between 100 – 200 fmol was required for end-prep, native barcode ligation and adaptor ligation. The final library was quantified with a Qubit® 2.0 Fluorometer to ensure that a final concentration of between 3 – 20 fmol was available. Libraries with higher concentrations were diluted with nuclease free water to ensure that they measured within the required range in a final volume of 5 µL.

The Flongle flow cell was checked, prior to sequencing initiation, to ensure that a minimum of 50 nanopores were available as recommended by ONT and to ensure sufficient sequencing quality. The library was prepared for sequencing by adding Sequencing Buffer II and Loading Beads II and added into the sample port of the Flongle flow cell. Runs were stopped once sufficient coverage of approximately 10 000X was obtained for all samples. High-accuracy basecalling with Guppy (version: 5.1.13), ‘trim-barcodes’, Q-score ≥ 8, Length range: 0.8 – 1.5 kb and the fast5 and fastq output files were compressed. Samples for which the ligation step had failed in a prior run were repeated on the next run, hence a total of three libraries were generated and sequenced for the 54 *PR* and *RT* nested-PCR amplicons and one library was generated and sequenced containing the 24 *IN* nested-PCR amplicons.

### Nano-RECall pipeline

The software performs protein-aware sequence alignments relative to four reference subtypes of each HIV-1 subtype (A-H) and common recombinant (CRF01_AE and CRF02_AG) to select the optimum alignment (independently for “forward” and “reverse” sequences) and identify sequence “mixtures”. It automatically incorporates sample and batch resistance reports based on the Stanford HIV drug resistance database (Stanford HIVDB) as well as Quality Control metrics (phylogenetic reports, sequence similarity flags and APOBEC mutation flags, etc) which are currently recommended by the World Health Organization[9].

The Nano-RECall pipeline generated consensus .fasta sequences, using a minor variant threshold of 25%. As a quality check and to compare consensus sequence similarity between Sanger and ONT sequences phylogenetic trees were first generated in Geneious Prime (version 2021.1.1) with the Tamura-Nei neighbor joining method.

### Statistics

We assessed percentage similarity between the two sequencing platforms and the respective bioinformatics interpretation algorithms, RECall and NanoRECall, for ONT and Sanger sequences; confidence intervals for mean sequence concordance was assessed with non-parametric bootstrapping in R 4.1.2[10]; Overall nucleotide level agreement between ONT and Sanger consensus sequences was assessed with Cohen’s kappa statistics.

Consensus sequences were interpreted at codon-level with the Stanford HIVDB version 9.0[11]. Overall agreement was assessed as the proportion of Stanford HIVDB -included mutations, detected by both platforms of those that were detected by either sequencing platform- and pipeline combination.

## Results

### Sanger sequences

Of 54 *PR-RT* sequences 7 were excluded that did not meet RECall quality criteria. Another sample was excluded based on phylogenetic analysis showing a sample mislabeling. Two of the 24 *IN* Sanger sequences did not meet RECall quality criteria and another had insufficient ligation during the ONT library preparation. The final analysis therefore included 46 *PR-RT* sequences and 21 *IN* Sequences.

Before comparing the sequencing platforms, we assessed the relationship between ONT sequence coverage depth and nucleotide variance.

The ONT Flongle run time was set to 24 hours (standard) and runs stopped when at least 10 000 read-depth was achieved for each barcoded sample. However, it became apparent that provided redundant read-depth to detect nucleotide mixtures at a level equivalent to Sanger sequencing. The coefficient of variation (CV%) of the codon proportion plateaus at ∼6% after a minimum read-depth of 400 fold (Figure 1), which allowed us to limit the amount of sequences processed. Randomly processing replicates of 400 sequences showed limited variation in variant frequency (Figure 2) with the overall proportion agreeing with electropherogram height (example of sample HIVDR68 shown in Figure 2). Phylogenetic trees (supplementary Figures) show identical sequences or close agreement between Sanger and ONT consensus; for the integrase sequence run samples INT_32 and INT_4 were different amplicons form the same patient, which showed close agreement.

**Figure 1.**
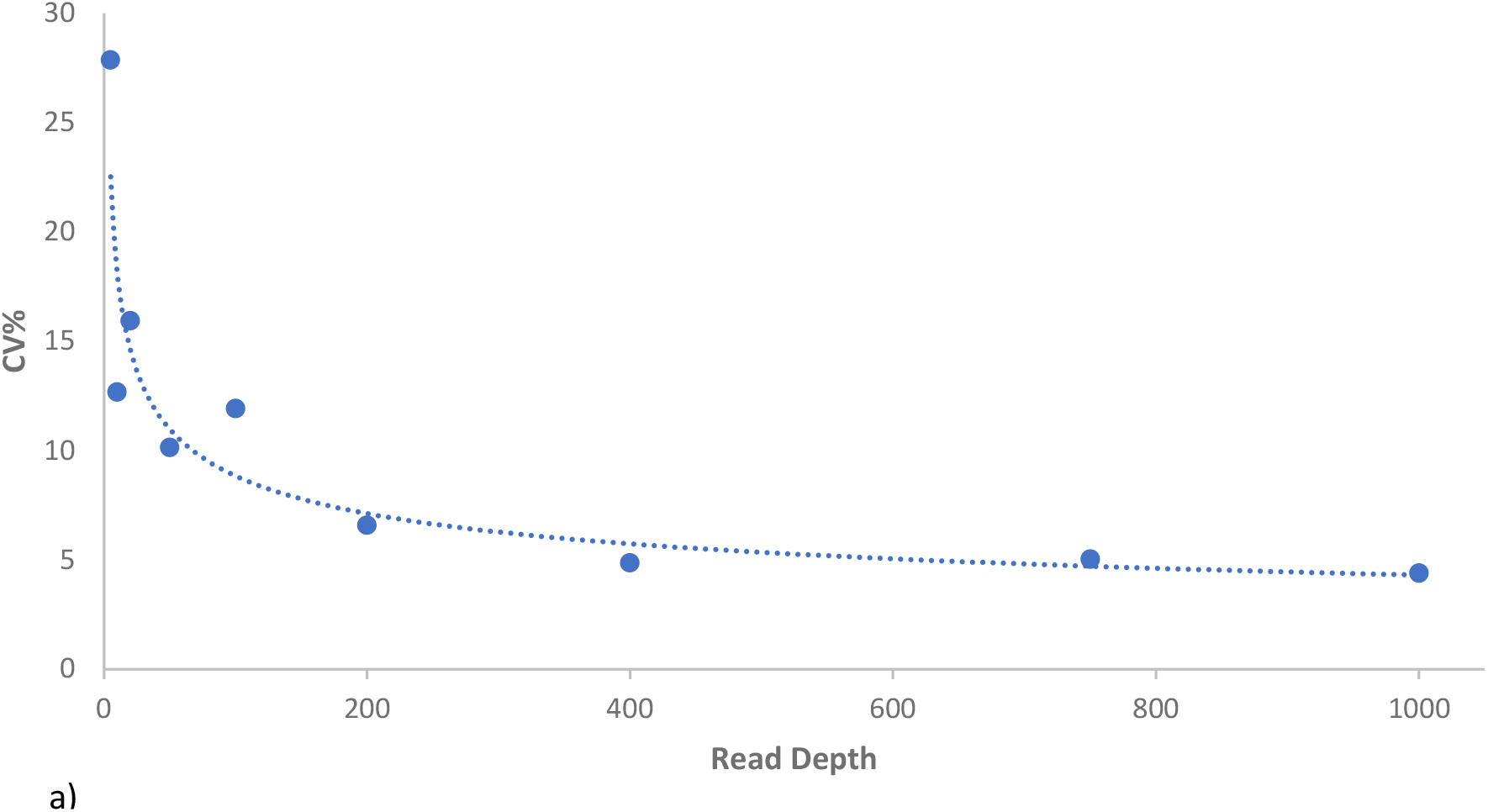
Relationship between sequence coverage depth and coefficient of variation. The coefficient of variation (CV%) of codon proportion is shown for varying sequence depth at common drug resistance mutations such as K103N.

**Figure 2.**
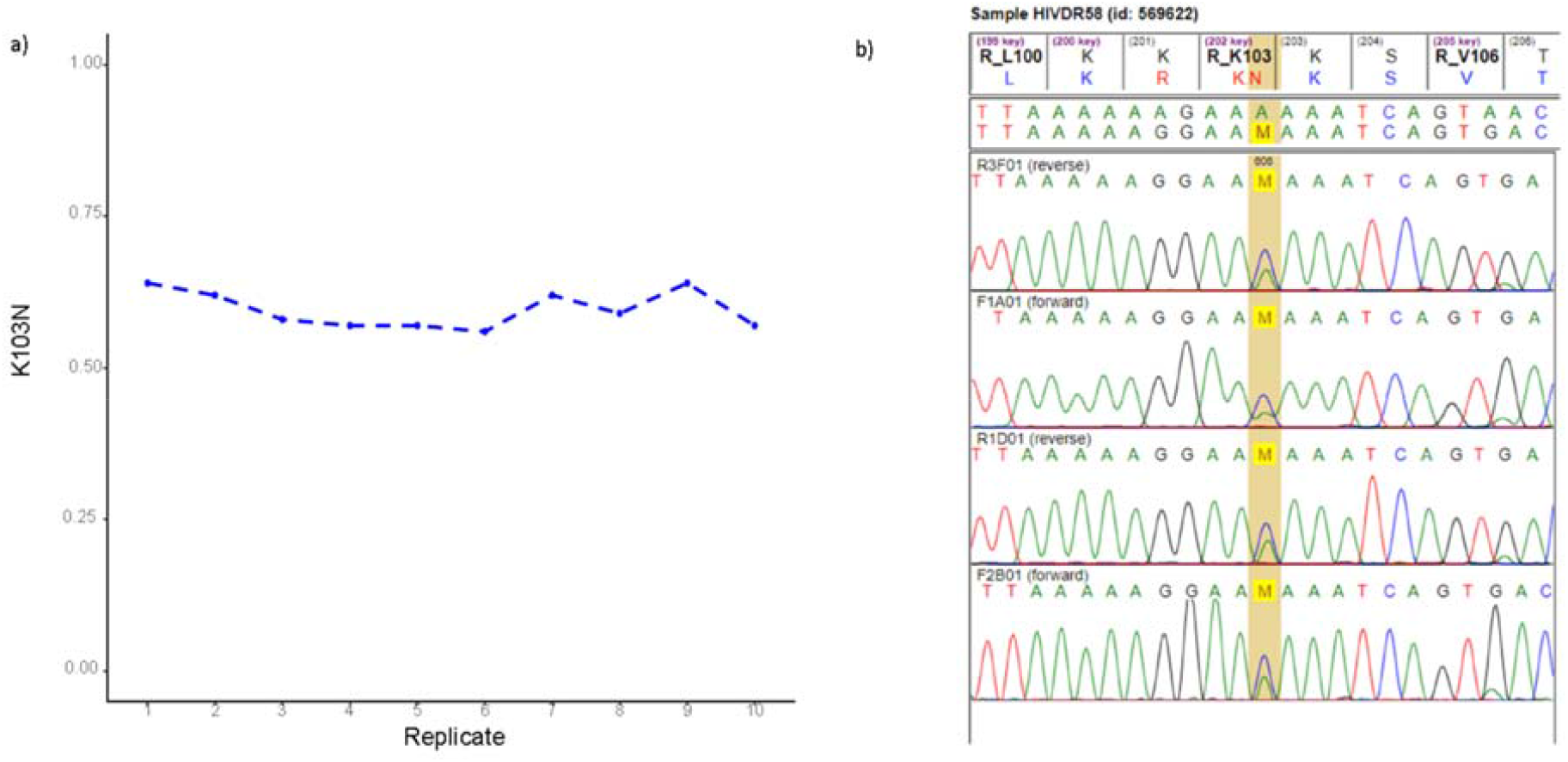
Variation in variant proportion for replicates of 400 reads. **a)** Variation in the proportion of K103N in replicates of 400 reads is shown for sample HIVDR58. b) The relative Sanger electropherogram peak high of cytosine (C) in AAC, encoding asparagine (N) is 69%. relative to all (including the wildtype base AAA, encoding wildtype lysine (K)).

For the 46 *PR-RT* and 21 *IN* amplicons included in the final analysis, Nano-RECall consensus sequences [GenBank accession numbers: ON720593 - ON720726] and RECall Sanger consensus sequences were compared on nucleotide level and with reference to drug resistance interpretation.

For *protease* and *reverse transcriptase* sequences the average consensus sequence similarity was 99.3% (95% CI: 99.1-99.34%), for *integrase* sequences: 99.6% (95% CI: 99.4-99.7%).

Agreement in consensus sequence bases are shown in Figure 3. There was an almost perfect overall agreement; Cohen’s kappa: 0.992 (95% CI: 0.991-0.993)[12].

**Figure 3.**
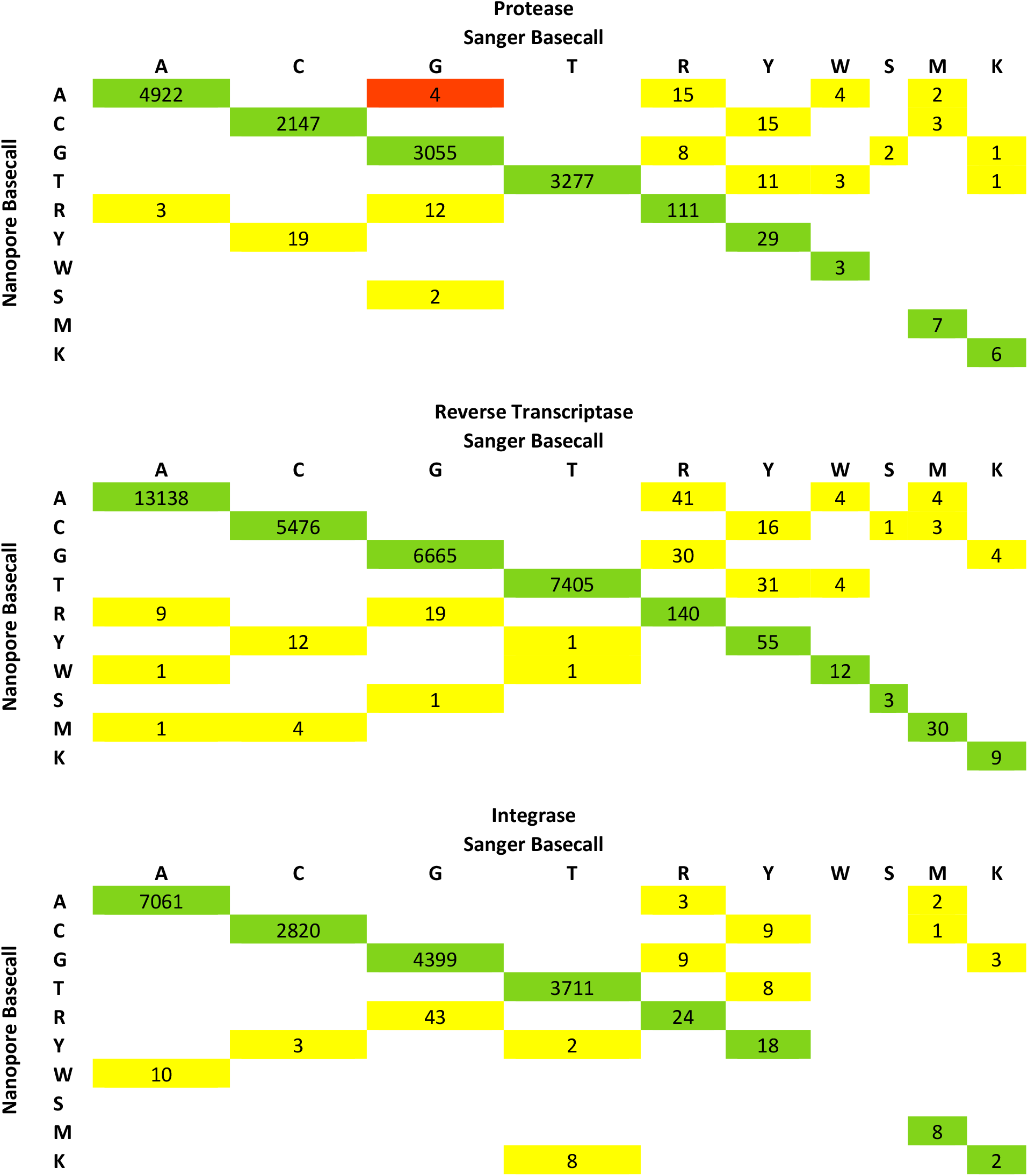
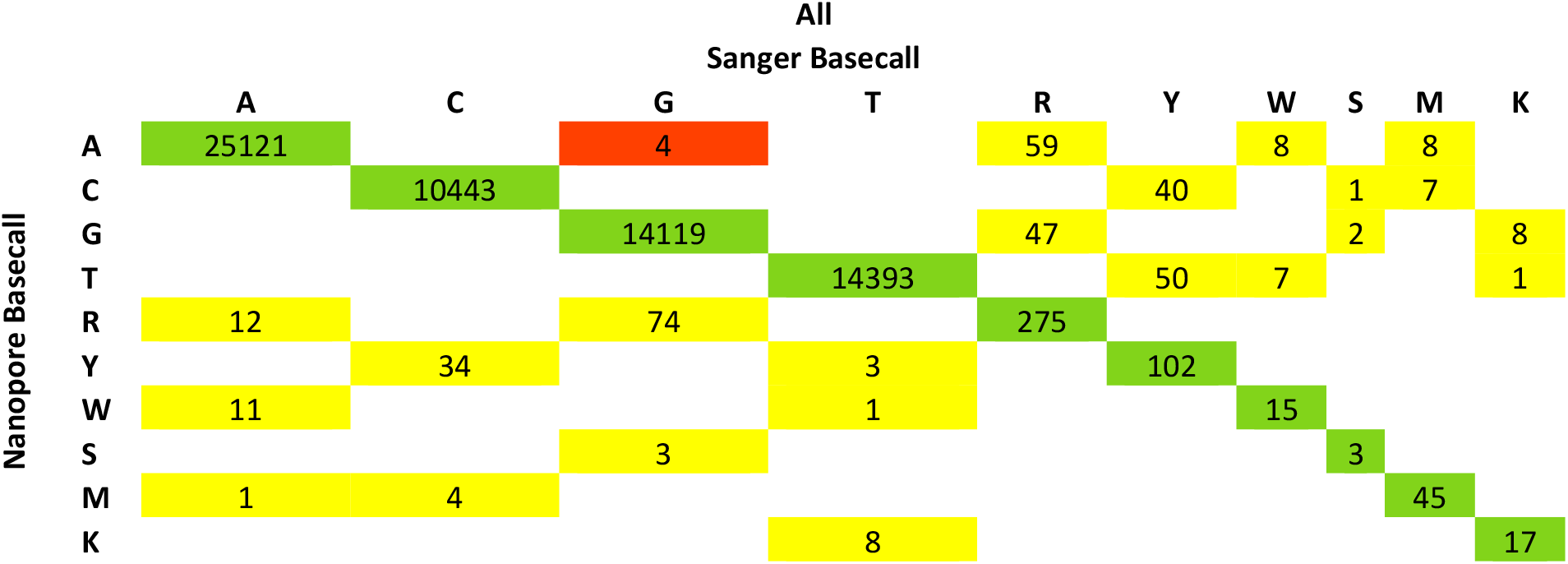
Agreement in base position for ONT and Sanger Sequencing. Agreement across all sequenced bases for *protease, reverse transcriptase, integrase* and “all positions” are shown. Green cells indicate agreement, yellow cells indicate partial agreement – mixed bases vs single bases- and red cells bases that were discordant between ONT Nano-RECall and Sanger RECall consensus sequences.

There was an overall good agreement in drug resistance interpretation between Nano-RECall and Sanger based drug resistance interpretation for either protease (Table 1.1; n=46), reverse transcriptase (Table 1.2; n=46) and for integrase (Table 1.3; n=21). In total there were 245 Stanford HIVDB mutations from the 46 cases with *PR* and *RT* sequences on either platform, with 241 mutations detected with ONT (of which 45 (18.7%) had codon mixtures detected) and 242 with Sanger (of which 55 (22.7%) had codon mixtures detected). Of the 245 mutations detected by either platform 238 (97.1%) were detected by both. With reference to protease inhibitor drug resistance mutations (Table 1.1) there were only discordance between codon-mixtures and single codons in 4 cases, 3 with reference to minor protease inhibitor drug resistance mutations (DR7, DR31 and DR33; all 3 detected by Sanger and not ONT) and one with reference to a major mutation (DR35; mixtures detected by Sanger and not ONT); hence none affected drug resistance level scores. With reference to NRTI drug resistance mutations, in one case (DR22) there was an accessory mutation, V75VM mixture detected by Sanger whereas the mutation was not detected by ONT, but this did not affect drug resistance level, another non-polymorphic mutation mixture D67DG (sample DR61), detected by Sanger but not ONT, increased resistance to AZT from intermediate to high. Three other sequences had partial disagreement with reference to NRTI resistance, DR35 and DR57 had mixtures detected by Sanger with single codons by ONT and DR58 a codon mixture by ONT and single codon by Sanger sequencing.

**Table 1.1:**
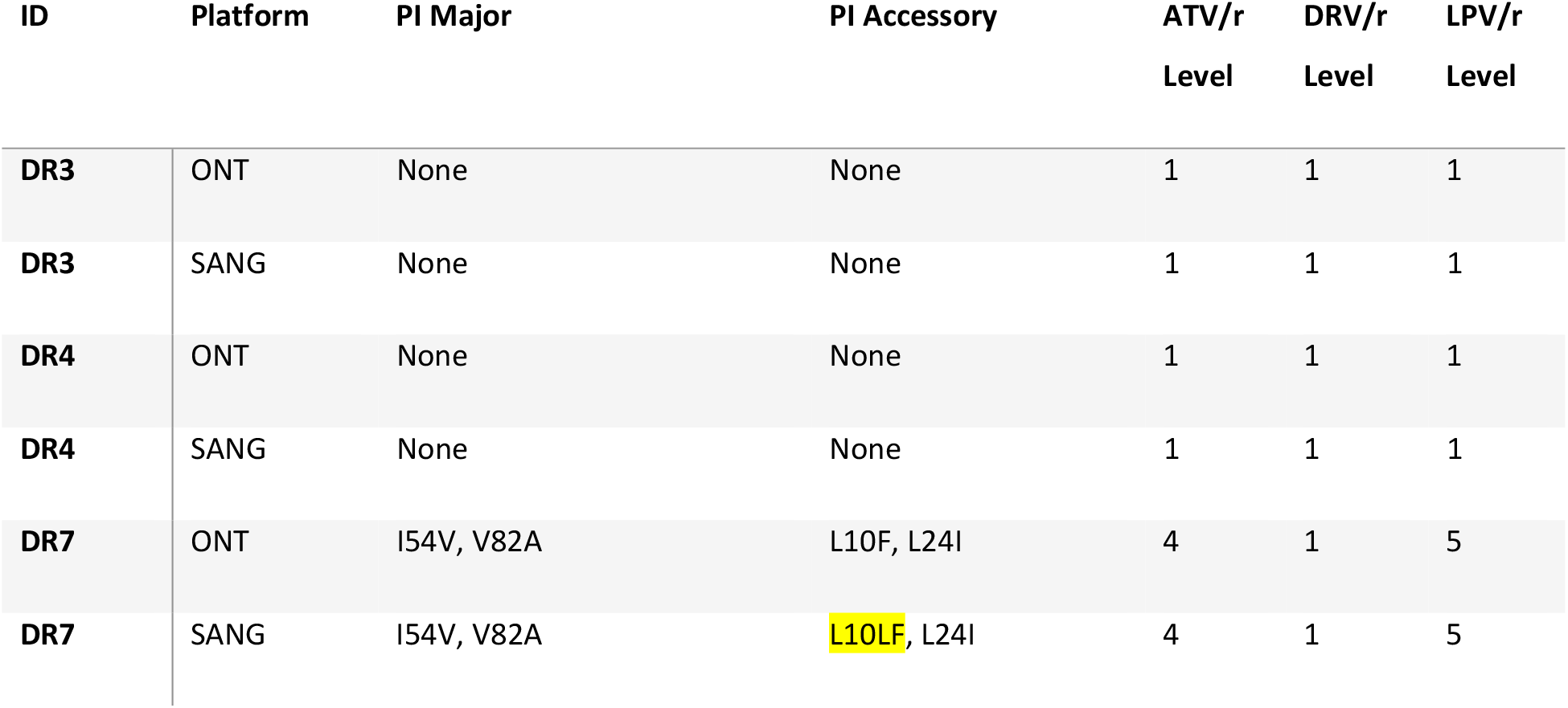
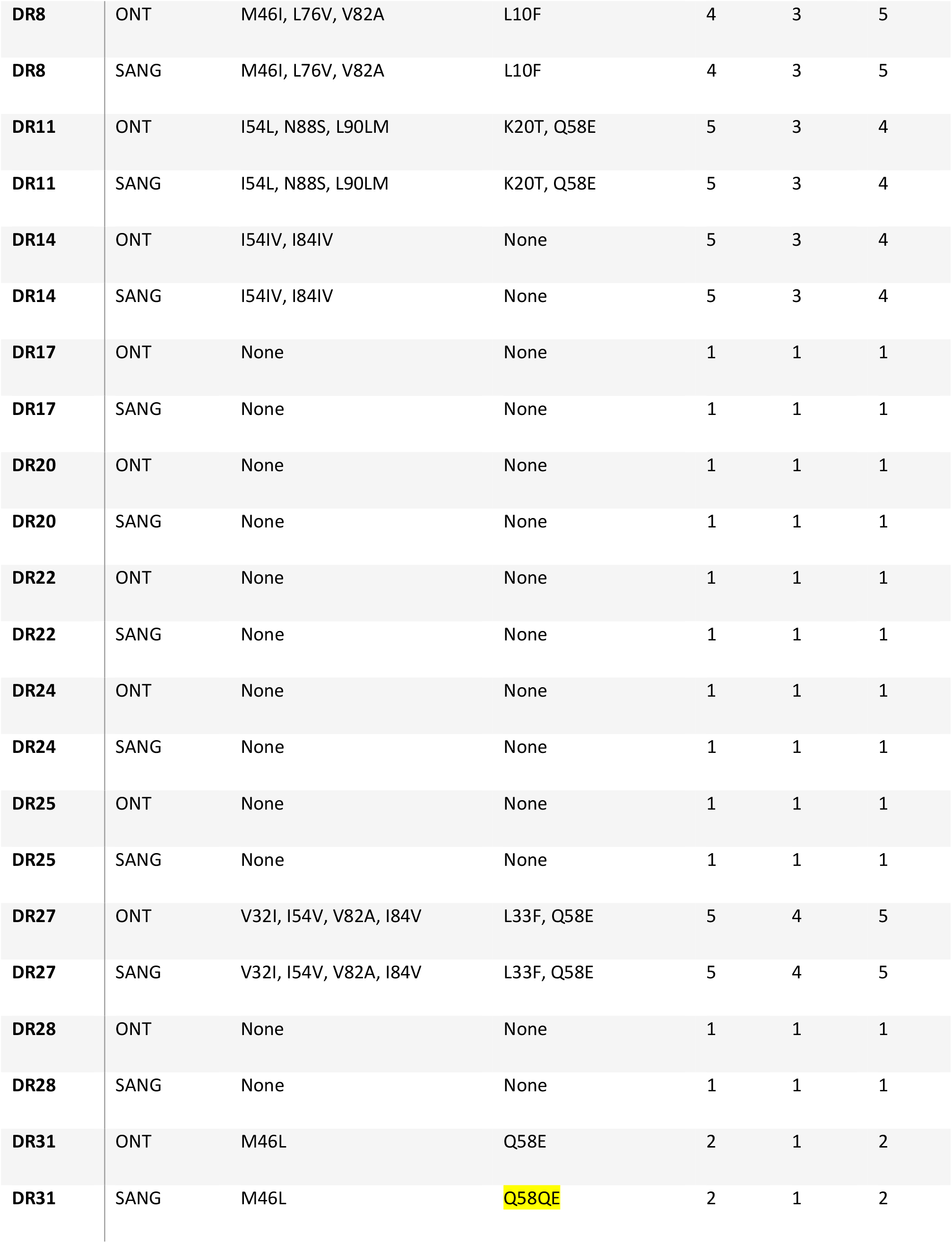
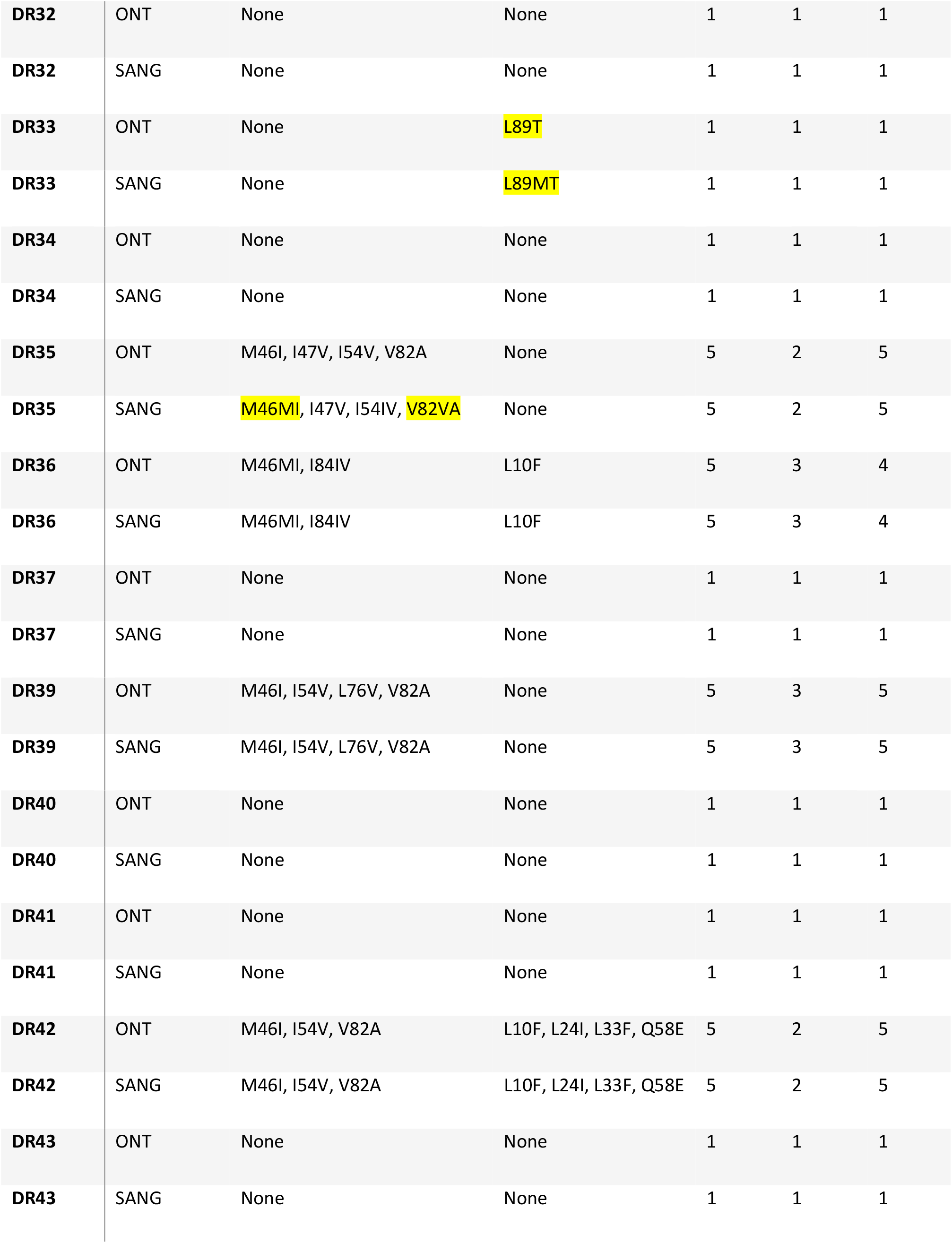
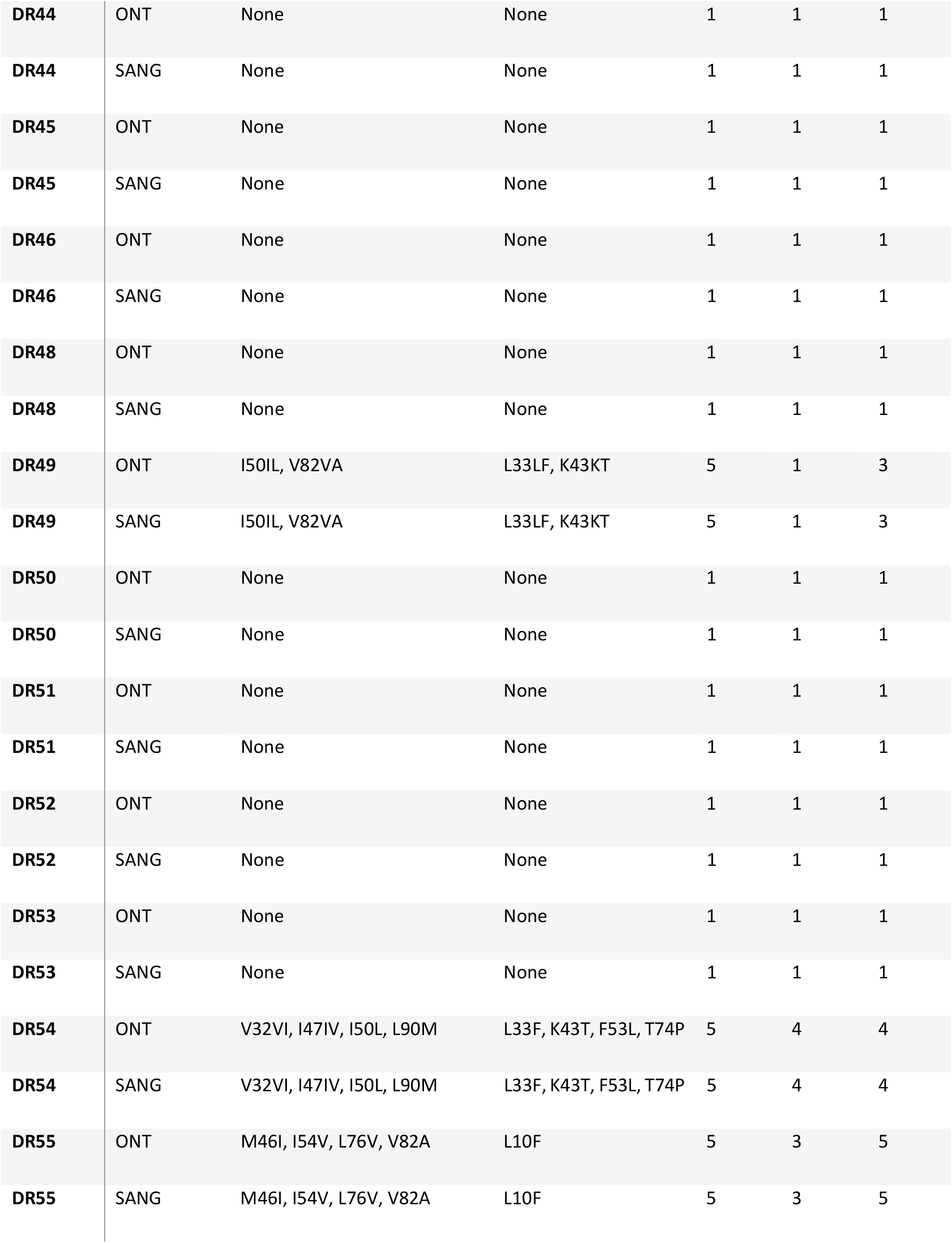
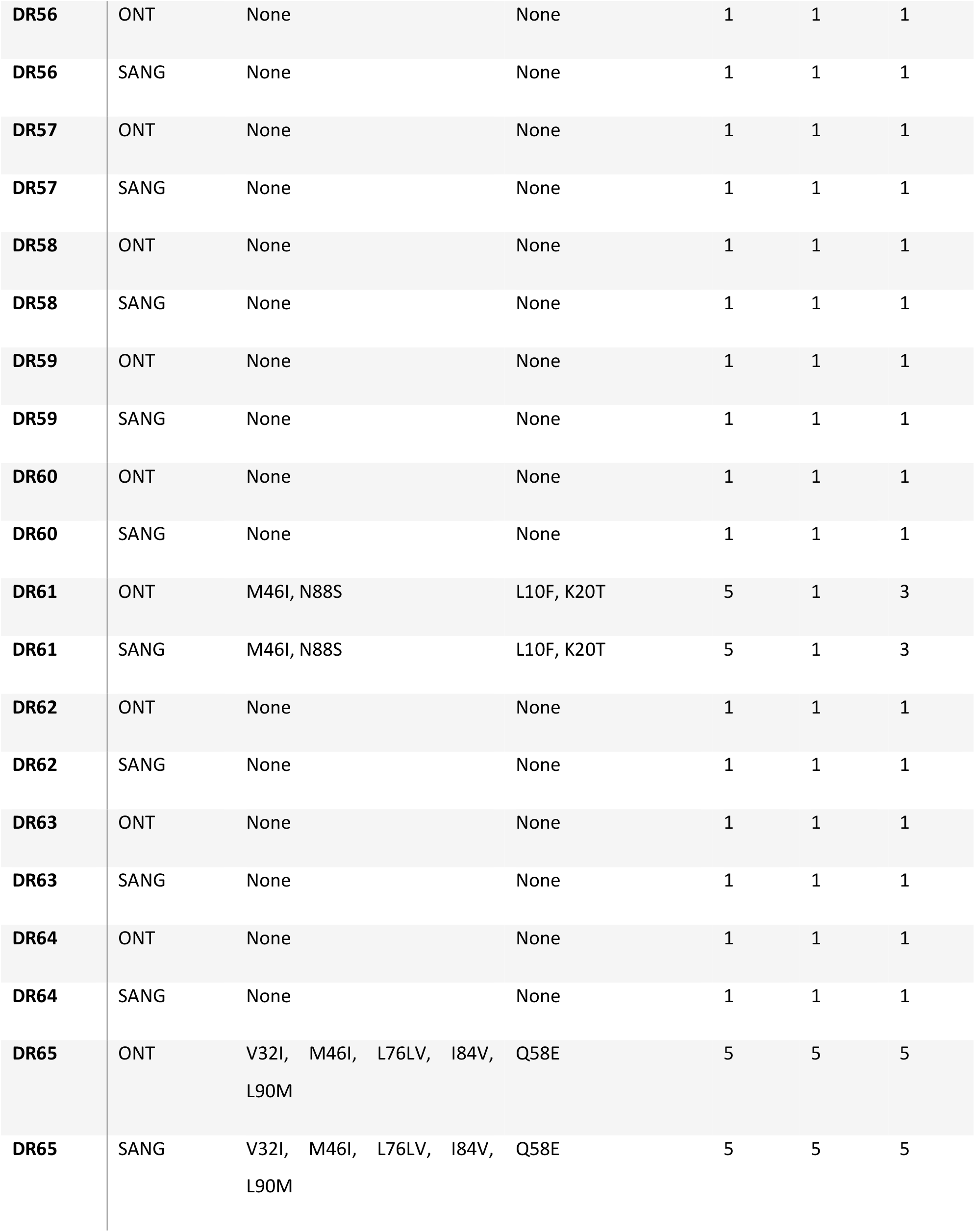
Stanford HIV DB drug resistance interpretation of 46 protease sequences that had ONT and Sanger sequencing respectively.

**Table 1.2:**
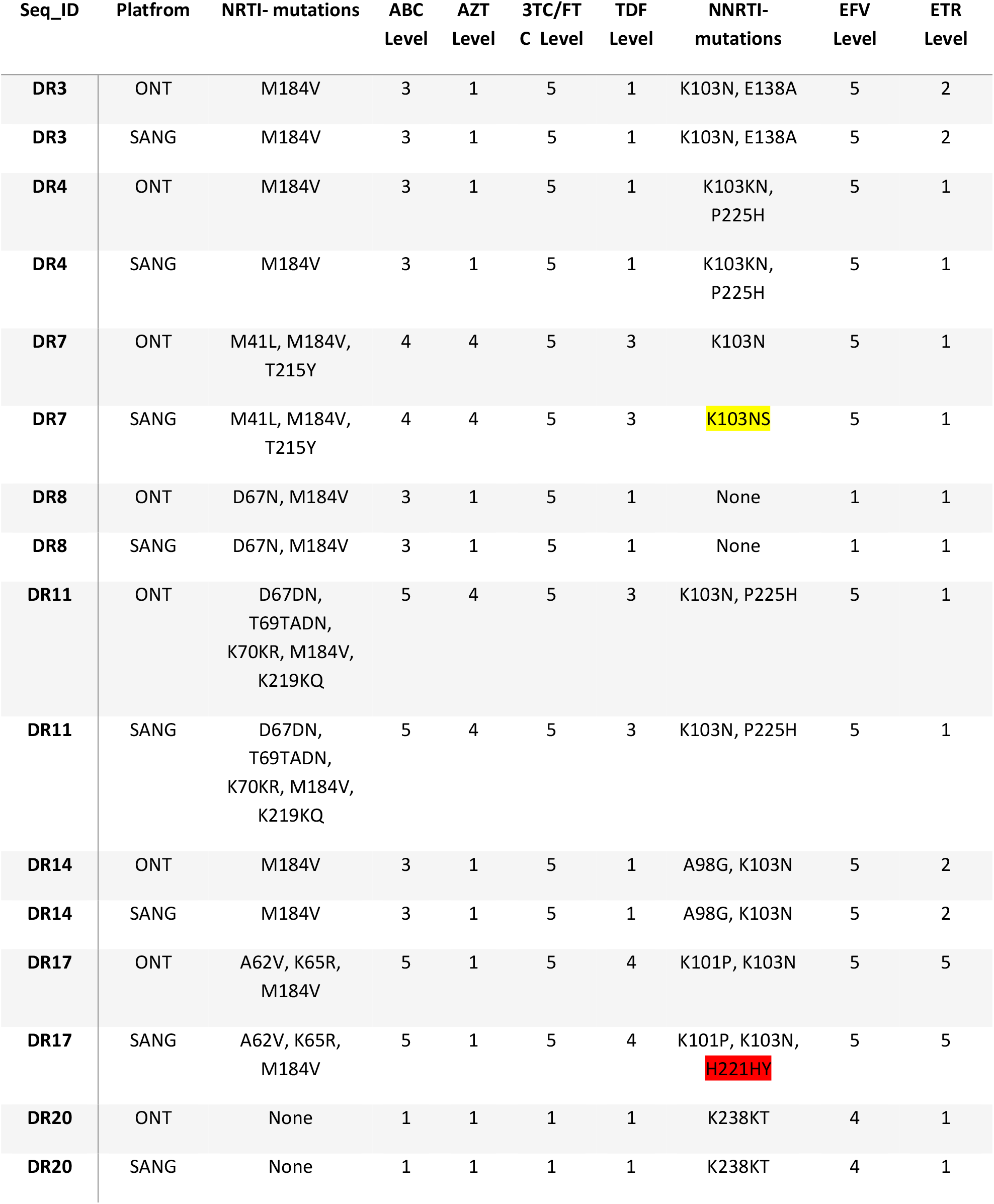
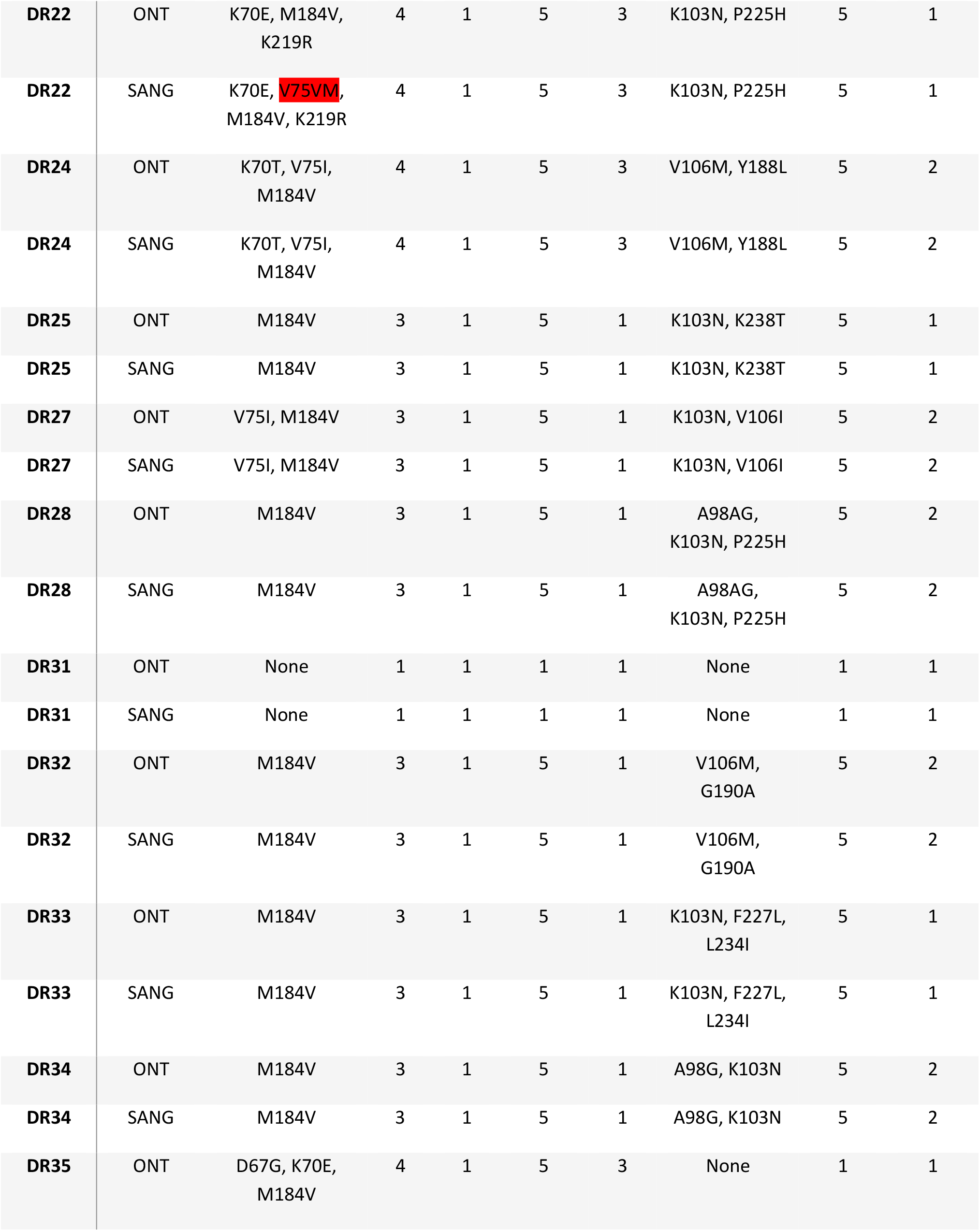
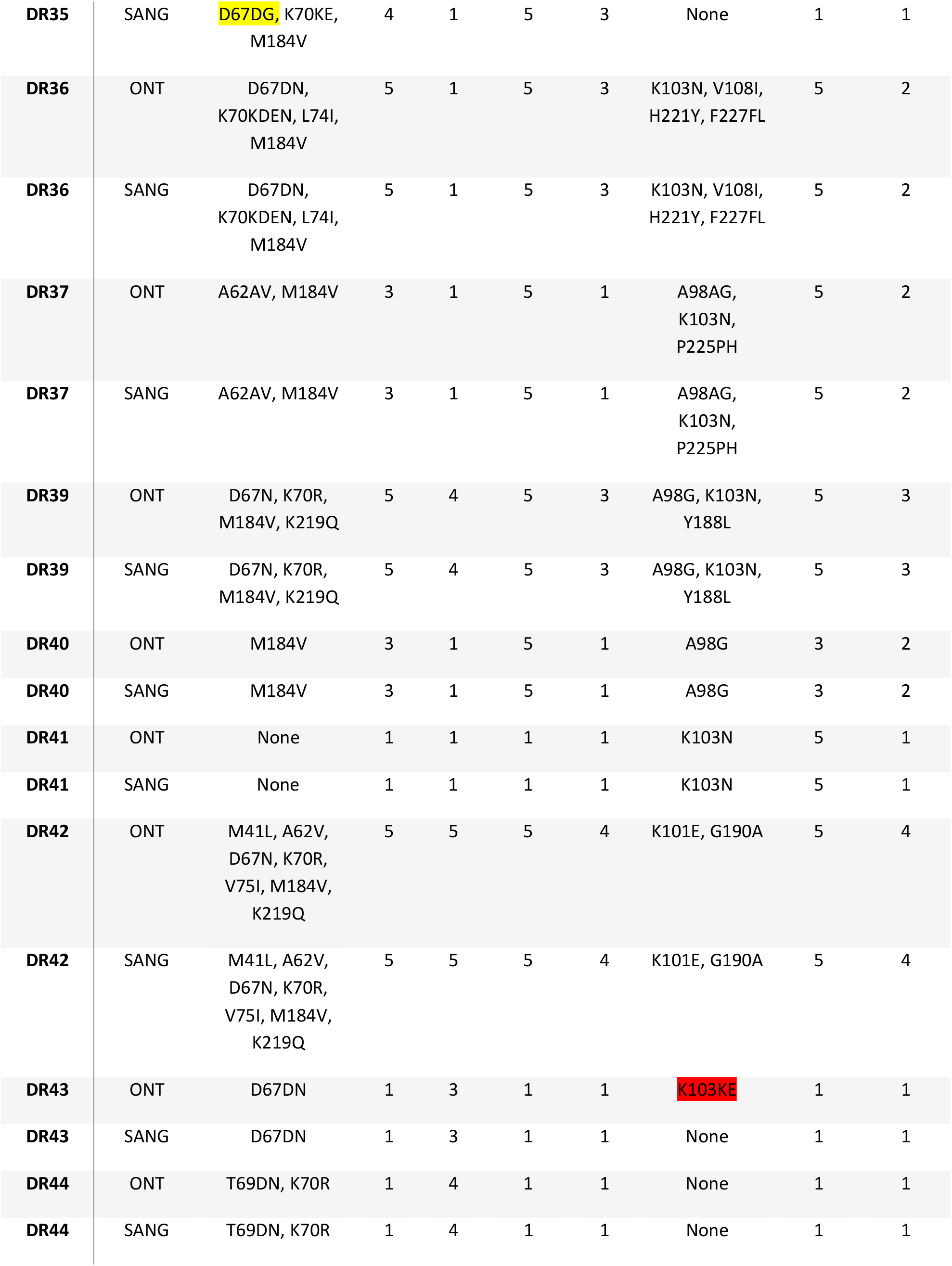
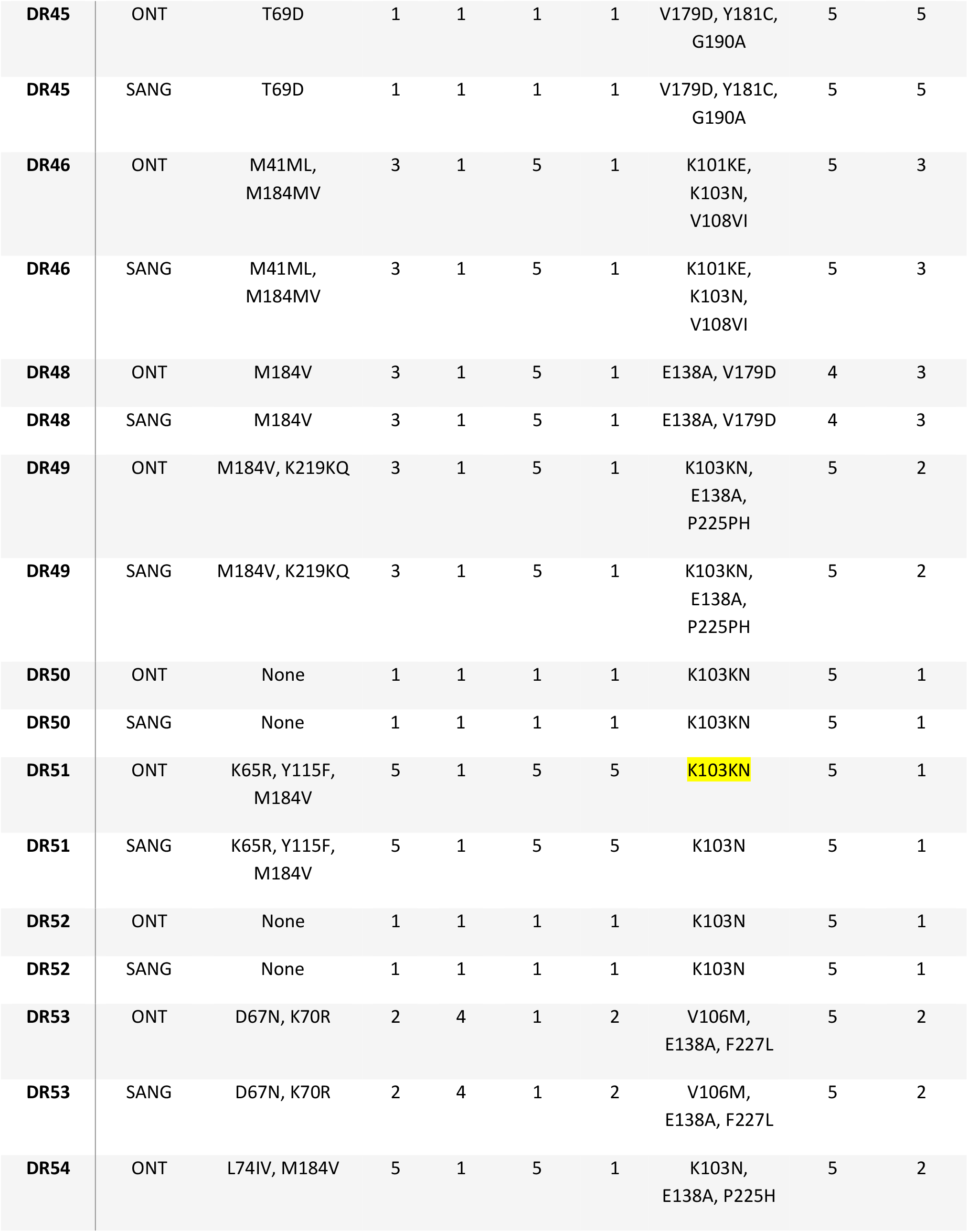
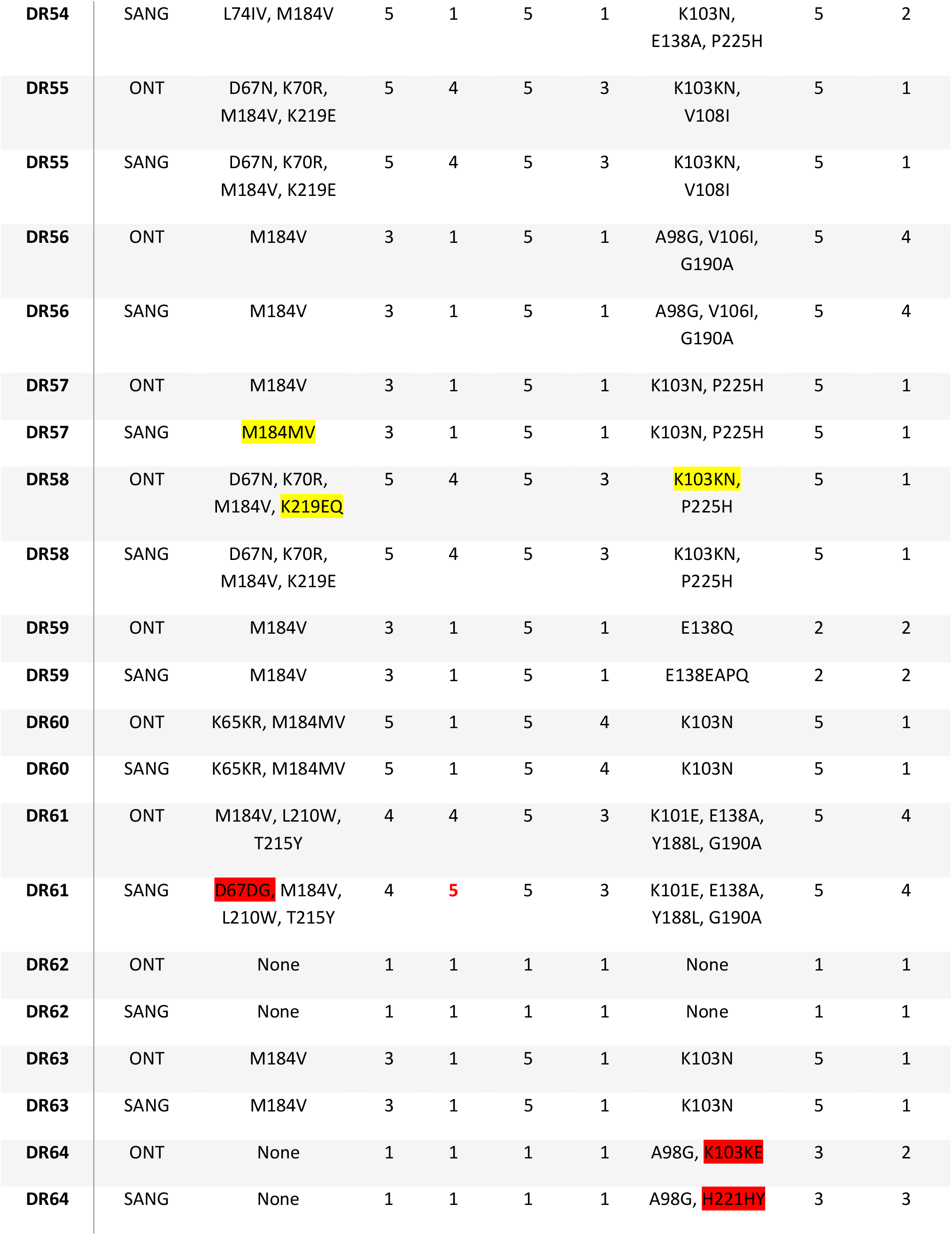
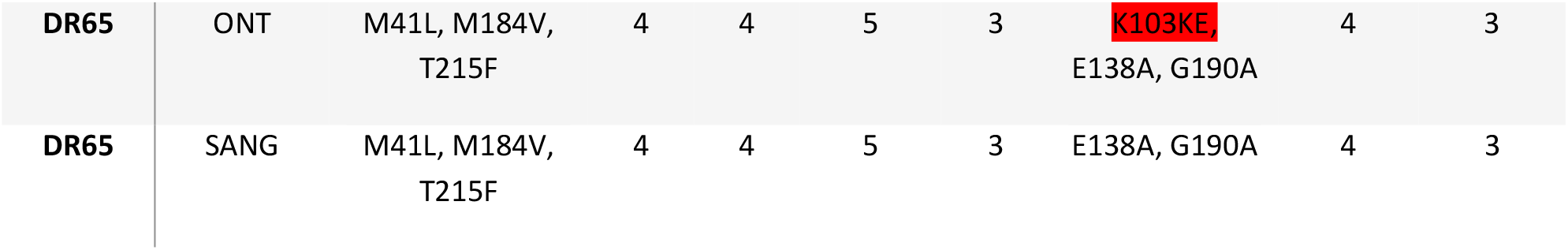
Stanford HIV DB drug resistance interpretation of 46 reverse transcriptase sequences that had ONT and Sanger sequencing respectively.

| Seq_ID | Platform | NRTI- mutations | ABC Level | AZT Level | 3TC/FT C Level | TDF Level | NNRTI- mutations | EFV Level | ETR Level |
| --- | --- | --- | --- | --- | --- | --- | --- | --- | --- |
| DR3 | ONT | M184V | 3 | 1 | 5 | 1 | K103N, E138A | 5 | 2 |
| DR3 | SANG | M184V | 3 | 1 | 5 | 1 | K103N, E138A | 5 | 2 |
| DR4 | ONT | M184V | 3 | 1 | 5 | 1 | K103KN, P225H | 5 | 1 |
| DR4 | SANG | M184V | 3 | 1 | 5 | 1 | K103KN, P225H | 5 | 1 |
| DR7 | ONT | M41L, M184V, T215Y | 4 | 4 | 5 | 3 | K103N | 5 | 1 |
| DR7 | SANG | M41L, M184V, T215Y | 4 | 4 | 5 | 3 | K103NS | 5 | 1 |
| DR8 | ONT | D67N, M184V | 3 | 1 | 5 | 1 | None | 1 | 1 |
| DR8 | SANG | D67N, M184V | 3 | 1 | 5 | 1 | None | 1 | 1 |
| DR11 | ONT | D67DN, T69TADN, K70KR, M184V, K219KQ | 5 | 4 | 5 | 3 | K103N, P225H | 5 | 1 |
| DR11 | SANG | D67DN, T69TADN, K70KR, M184V, K219KQ | 5 | 4 | 5 | 3 | K103N, P225H | 5 | 1 |
| DR14 | ONT | M184V | 3 | 1 | 5 | 1 | A98G, K103N | 5 | 2 |
| DR14 | SANG | M184V | 3 | 1 | 5 | 1 | A98G, K103N | 5 | 2 |
| DR17 | ONT | A62V, K65R, M184V | 5 | 1 | 5 | 4 | K101P, K103N | 5 | 5 |
| DR17 | SANG | A62V, K65R, M184V | 5 | 1 | 5 | 4 | K101P, K103N, H221HY | 5 | 5 |
| DR20 | ONT | None | 1 | 1 | 1 | 1 | K238KT | 4 | 1 |
| DR20 | SANG | None | 1 | 1 | 1 | 1 | K238KT | 4 | 1 |
| DR22 | ONT | K70E, M184V,<br>K219R | 4 | 1 | 5 | 3 | K103N, P225H | 5 | 1 |
| DR22 | SANG | K70E, V75VM,<br>M184V, K219R | 4 | 1 | 5 | 3 | K103N, P225H | 5 | 1 |
| DR24 | ONT | K70T, V75I,<br>M184V | 4 | 1 | 5 | 3 | V106M, Y188L | 5 | 2 |
| DR24 | SANG | K70T, V75I,<br>M184V | 4 | 1 | 5 | 3 | V106M, Y188L | 5 | 2 |
| DR25 | ONT | M184V | 3 | 1 | 5 | 1 | K103N, K238T | 5 | 1 |
| DR25 | SANG | M184V | 3 | 1 | 5 | 1 | K103N, K238T | 5 | 1 |
| DR27 | ONT | V75I, M184V | 3 | 1 | 5 | 1 | K103N, V106I | 5 | 2 |
| DR27 | SANG | V75I, M184V | 3 | 1 | 5 | 1 | K103N, V106I | 5 | 2 |
| DR28 | ONT | M184V | 3 | 1 | 5 | 1 | A98AG,<br>K103N, P225H | 5 | 2 |
| DR28 | SANG | M184V | 3 | 1 | 5 | 1 | A98AG,<br>K103N, P225H | 5 | 2 |
| DR31 | ONT | None | 1 | 1 | 1 | 1 | None | 1 | 1 |
| DR31 | SANG | None | 1 | 1 | 1 | 1 | None | 1 | 1 |
| DR32 | ONT | M184V | 3 | 1 | 5 | 1 | V106M,<br>G190A | 5 | 2 |
| DR32 | SANG | M184V | 3 | 1 | 5 | 1 | V106M,<br>G190A | 5 | 2 |
| DR33 | ONT | M184V | 3 | 1 | 5 | 1 | K103N, F227L,<br>L234I | 5 | 1 |
| DR33 | SANG | M184V | 3 | 1 | 5 | 1 | K103N, F227L,<br>L234I | 5 | 1 |
| DR34 | ONT | M184V | 3 | 1 | 5 | 1 | A98G, K103N | 5 | 2 |
| DR34 | SANG | M184V | 3 | 1 | 5 | 1 | A98G, K103N | 5 | 2 |
| DR35 | ONT | D67G, K70E,<br>M184V | 4 | 1 | 5 | 3 | None | 1 | 1 |
| DR35 | SANG | D67DG, K70KE,<br>M184V | 4 | 1 | 5 | 3 | None | 1 | 1 |
| DR36 | ONT | D67DN,<br>K70KDEN, L74I,<br>M184V | 5 | 1 | 5 | 3 | K103N, V108I,<br>H221Y, F227FL | 5 | 2 |
| DR36 | SANG | D67DN,<br>K70KDEN, L74I,<br>M184V | 5 | 1 | 5 | 3 | K103N, V108I,<br>H221Y, F227FL | 5 | 2 |
| DR37 | ONT | A62AV, M184V | 3 | 1 | 5 | 1 | A98AG,<br>K103N,<br>P225PH | 5 | 2 |
| DR37 | SANG | A62AV, M184V | 3 | 1 | 5 | 1 | A98AG,<br>K103N,<br>P225PH | 5 | 2 |
| DR39 | ONT | D67N, K70R,<br>M184V, K219Q | 5 | 4 | 5 | 3 | A98G, K103N,<br>Y188L | 5 | 3 |
| DR39 | SANG | D67N, K70R,<br>M184V, K219Q | 5 | 4 | 5 | 3 | A98G, K103N,<br>Y188L | 5 | 3 |
| DR40 | ONT | M184V | 3 | 1 | 5 | 1 | A98G | 3 | 2 |
| DR40 | SANG | M184V | 3 | 1 | 5 | 1 | A98G | 3 | 2 |
| DR41 | ONT | None | 1 | 1 | 1 | 1 | K103N | 5 | 1 |
| DR41 | SANG | None | 1 | 1 | 1 | 1 | K103N | 5 | 1 |
| DR42 | ONT | M41L, A62V,<br>D67N, K70R,<br>V75I, M184V,<br>K219Q | 5 | 5 | 5 | 4 | K101E, G190A | 5 | 4 |
| DR42 | SANG | M41L, A62V,<br>D67N, K70R,<br>V75I, M184V,<br>K219Q | 5 | 5 | 5 | 4 | K101E, G190A | 5 | 4 |
| DR43 | ONT | D67DN | 1 | 3 | 1 | 1 | K103KE | 1 | 1 |
| DR43 | SANG | D67DN | 1 | 3 | 1 | 1 | None | 1 | 1 |
| DR44 | ONT | T69DN, K70R | 1 | 4 | 1 | 1 | None | 1 | 1 |
| DR44 | SANG | T69DN, K70R | 1 | 4 | 1 | 1 | None | 1 | 1 |
| DR45 | ONT | T69D | 1 | 1 | 1 | 1 | V179D, Y181C,<br>G190A | 5 | 5 |
| DR45 | SANG | T69D | 1 | 1 | 1 | 1 | V179D, Y181C,<br>G190A | 5 | 5 |
| DR46 | ONT | M41ML,<br>M184MV | 3 | 1 | 5 | 1 | K101KE,<br>K103N,<br>V108VI | 5 | 3 |
| DR46 | SANG | M41ML,<br>M184MV | 3 | 1 | 5 | 1 | K101KE,<br>K103N,<br>V108VI | 5 | 3 |
| DR48 | ONT | M184V | 3 | 1 | 5 | 1 | E138A, V179D | 4 | 3 |
| DR48 | SANG | M184V | 3 | 1 | 5 | 1 | E138A, V179D | 4 | 3 |
| DR49 | ONT | M184V, K219KQ | 3 | 1 | 5 | 1 | K103KN,<br>E138A,<br>P225PH | 5 | 2 |
| DR49 | SANG | M184V, K219KQ | 3 | 1 | 5 | 1 | K103KN,<br>E138A,<br>P225PH | 5 | 2 |
| DR50 | ONT | None | 1 | 1 | 1 | 1 | K103KN | 5 | 1 |
| DR50 | SANG | None | 1 | 1 | 1 | 1 | K103KN | 5 | 1 |
| DR51 | ONT | K65R, Y115F,<br>M184V | 5 | 1 | 5 | 5 | K103KN | 5 | 1 |
| DR51 | SANG | K65R, Y115F,<br>M184V | 5 | 1 | 5 | 5 | K103N | 5 | 1 |
| DR52 | ONT | None | 1 | 1 | 1 | 1 | K103N | 5 | 1 |
| DR52 | SANG | None | 1 | 1 | 1 | 1 | K103N | 5 | 1 |
| DR53 | ONT | D67N, K70R | 2 | 4 | 1 | 2 | V106M,<br>E138A, F227L | 5 | 2 |
| DR53 | SANG | D67N, K70R | 2 | 4 | 1 | 2 | V106M,<br>E138A, F227L | 5 | 2 |
| DR54 | ONT | L74IV, M184V | 5 | 1 | 5 | 1 | K103N,<br>E138A, P225H | 5 | 2 |
| DR54 | SANG | L74IV, M184V | 5 | 1 | 5 | 1 | K103N,<br>E138A, P225H | 5 | 2 |
| DR55 | ONT | D67N, K70R,<br>M184V, K219E | 5 | 4 | 5 | 3 | K103KN,<br>V108I | 5 | 1 |
| DR55 | SANG | D67N, K70R,<br>M184V, K219E | 5 | 4 | 5 | 3 | K103KN,<br>V108I | 5 | 1 |
| DR56 | ONT | M184V | 3 | 1 | 5 | 1 | A98G, V106I,<br>G190A | 5 | 4 |
| DR56 | SANG | M184V | 3 | 1 | 5 | 1 | A98G, V106I,<br>G190A | 5 | 4 |
| DR57 | ONT | M184V | 3 | 1 | 5 | 1 | K103N, P225H | 5 | 1 |
| DR57 | SANG | M184MV | 3 | 1 | 5 | 1 | K103N, P225H | 5 | 1 |
| DR58 | ONT | D67N, K70R,<br>M184V, K219EQ | 5 | 4 | 5 | 3 | K103KN,<br>P225H | 5 | 1 |
| DR58 | SANG | D67N, K70R,<br>M184V, K219E | 5 | 4 | 5 | 3 | K103KN,<br>P225H | 5 | 1 |
| DR59 | ONT | M184V | 3 | 1 | 5 | 1 | E138Q | 2 | 2 |
| DR59 | SANG | M184V | 3 | 1 | 5 | 1 | E138EAPQ | 2 | 2 |
| DR60 | ONT | K65KR, M184MV | 5 | 1 | 5 | 4 | K103N | 5 | 1 |
| DR60 | SANG | K65KR, M184MV | 5 | 1 | 5 | 4 | K103N | 5 | 1 |
| DR61 | ONT | M184V, L210W,<br>T215Y | 4 | 4 | 5 | 3 | K101E, E138A,<br>Y188L, G190A | 5 | 4 |
| DR61 | SANG | D67DG, M184V,<br>L210W, T215Y | 4 | 5 | 5 | 3 | K101E, E138A,<br>Y188L, G190A | 5 | 4 |
| DR62 | ONT | None | 1 | 1 | 1 | 1 | None | 1 | 1 |
| DR62 | SANG | None | 1 | 1 | 1 | 1 | None | 1 | 1 |
| DR63 | ONT | M184V | 3 | 1 | 5 | 1 | K103N | 5 | 1 |
| DR63 | SANG | M184V | 3 | 1 | 5 | 1 | K103N | 5 | 1 |
| DR64 | ONT | None | 1 | 1 | 1 | 1 | A98G, K103KE | 3 | 2 |
| DR64 | SANG | None | 1 | 1 | 1 | 1 | A98G, H221HY | 3 | 3 |
| DR65 | ONT | M41L, M184V,<br>T215F | 4 | 4 | 5 | 3 | K103KE<br>E138A, G190A | 4 | 3 |
| DR65 | SANG | M41L, M184V,<br>T215F | 4 | 4 | 5 | 3 | E138A, G190A | 4 | 3 |

**Table 1.3:** Stanford HIV DB drug resistance interpretation of 21 integrase sequences that had ONT and Sanger sequencing respectively. **Legend of Table 1**: The same PCR amplicons were respectively sequenced on Oxford Nanopore Technology Flongle flow cells (ONT platform) or by automated Sanger Sequencing (SANG platform). Drug resistance levels are shown for some antiretroviral drugs commonly used in developing countries using the Stanford Drug Resistance Database 5 level score: 1- susceptible, 2- potential low-level resistance, 3 – low-level resistance, 4- intermediate level resistance and 5- high level resistance. Protease inhibitors (PIs) included here are atazanavir/ritonavir (ATZ/r), lopinavir/ritonavir (LPV/r) and darunavir/ritonavir (DRV/r). Nucleos(t)ide reverse transcriptase inhibitors (NRTIs) are abacavir, zidovudine (AZT), lamivudine (3TC), emtricitabine (FTC) and tenofovir disoproxil fumarate (TDF). Non-nucleoside reverse transcriptase inhibitors (NNRTIs) included here are efavirenz (EFV) and etravirine (ETR). Discordant cases are highlighted in yellow when there was partial agreement – codon mixture and single codon and in red when there was a drug resistance mutation detected on the one platform but not on the other.

| ID | Platform | INSTI Major | INSTI Accessory | DTG<br>Level | RAL<br>Level |
| --- | --- | --- | --- | --- | --- |
| INT_1 | ONT | None | H51Y | 2 | 3 |
| INT_1 | SANG | None | H51Y | 2 | 3 |
| INT_2 | ONT | N155H,<br>R263K | None | 5 | 5 |
| INT_2 | SANG | N155H,<br>R263K | None | 5 | 5 |
| INT_3 | ONT | None | None | 1 | 1 |
| INT_3 | SANG | None | None | 1 | 1 |
| INT_4 | ONT | None | None | 1 | 1 |
| INT_4 | SANG | None | None | 1 | 1 |
| INT_5 | ONT | None | E157Q | 1 | 2 |
| INT_5 | SANG | None | E157Q | 1 | 2 |
| INT_6 | ONT | None | Q95K | 1 | 2 |
| INT_6 | SANG | None | Q95K | 1 | 2 |
| INT_9 | ONT | None | None | 1 | 1 |
| INT_9 | SANG | None | None | 1 | 1 |
| INT_10 | ONT | None | None | 1 | 1 |
| INT_10 | SANG | None | None | 1 | 1 |
| INT_11 | ONT | None | None | 1 | 1 |
| INT_11 | SANG | None | None | 1 | 1 |
| INT_13 | ONT | None | None | 1 | 1 |
| INT_13 | SANG | None | None | 1 | 1 |
| INT_14 | ONT | None | E157Q | 1 | 2 |
| INT_14 | SANG | None | E157Q | 1 | 2 |
| INT_17 | ONT | None | None | 1 | 1 |
| INT_17 | SANG | None | None | 1 | 1 |
| INT_18 | ONT | None | None | 1 | 1 |
| INT_18 | SANG | None | None | 1 | 1 |
| INT_19 | ONT | None | None | 1 | 1 |
| INT_19 | SANG | None | None | 1 | 1 |
| INT_20 | ONT | None | None | 1 | 1 |
| INT_20 | SANG | None | None | 1 | 1 |
| INT_21 | ONT | None | None | 1 | 1 |
| INT_21 | SANG | None | None | 1 | 1 |
| INT_22 | ONT | None | None | 1 | 1 |
| INT_22 | SANG | None | None | 1 | 1 |
| INT_23 | ONT | None | None | 1 | 1 |
| INT_23 | SANG | None | None | 1 | 1 |
| INT_27 | ONT | None | None | 1 | 1 |
| INT_27 | SANG | None | None | 1 | 1 |
| INT_31 | ONT | G118R,<br>E138K | None | 5 | 5 |
| INT_31 | SANG | G118R,<br>E138K | None | 5 | 5 |
| INT_32 | ONT | None | None | 1 | 1 |
| INT_32 | SANG | None | None | 1 | 1 |

With reference to NNRTI mutations, in two cases Sanger sequencing detected mixed codons vs single codons with ONT (samples DR7 and DR59), and in one case (sample DR51) ONT detected a mixed codon whereas Sanger detected a single codon. In one sample (DR17) Sanger detected H221HY, not detected by ONT and in two samples, DR43 and DR65 ONT detected K103E, not detected by Sanger, whereas in sample DR64 ONT detected K103KE whereas Sanger detected H221HY, not detected by the other platform in each case. Resistance interpretation for efavirenz (EFV) and etravirine (ETR) agreed in all cases except sample DR64 where the Sanger interpretation indicated low level ETR resistance, compared to potential low level ETR resistance with ONT.

There were no discordant drug resistance mutations among 16 Stanford HIVDB mutations from 21 IN sequences between ONT- and Sanger sequencing (Table 1.3).

## Discussion

Nano-RECall is software specifically designed for simple processing of HIV sequences from Oxford Nanopore devices, with a focus on drug resistance. Designed to be as similar for the user to the widely used “RECall” program as possible, the Ruby-based software is available as a freely distributed Windows modules or a web interface for those who prefer that approach. The Nano-RECall software is designed to be rapid and easy-to use while accommodating ONT homopolymer errors. Particular advantages of the software are that it is Windows-based and/or Web-based and have a similar interface to a currently widely used approach, in order to lower the learning curve for users. Using the Nano-RECall pipeline we found a high sequence similarity of >99% between ONT and Sanger sequencing for both *PR-RT* and *IN* sequencing.

The strength of this comparison was that we compared sequencing results using the same PCR amplicon. This eliminated the influence of PCR resampling error and PCR introduced mutations on the findings[13]. It therefore enabled us to assess the impact of the sequencing platforms, automated Sanger sequencing versus Oxford Nanopore Flongle sequencing coupled with their respective bioinformatics pipelines. The independent analysis of fastq data collected in the 5’ and 3’ directions provides a built-in quality check for each base called. This approach differed from generic approaches as it was specifically designed to correct for ONT homopolymer associated read error. Our study had several limitations. Our analysis was limited to high abundance drug resistance variants. The primary purpose was not to detect minor or low abundance variants but to investigate a Sanger sequencing alternative. Moreover, despite improvements in basecalling accuracy with new ONT chemistries, we remained concerned that homopolymer errors would result in the spurious detection of drug resistance mutations. We therefore set a threshold of 25% for ONT minor variants. This likely resulted in some of the discordant cases where Sanger sequencing detected codon mixtures but ONT did not. ONT has been improving their chemistry and basecalling algorithms resulting in a rapid reduction in read-error[14,15]. Our study was limited to samples from South Africa where HIV-1 subtype C predominates. In future Nano-RECall should be investigated using samples from other subtypes. We also did not compare ONT sequencing and Nano-RECall to high-depth next generation sequencing. In future, as the read accuracy of ONT improves further, one would be able to assess Nano-RECall using a lower variant detection threshold, but this would likely require processing more than 400 reads. As our study is a laboratory-based anonymized comparison we did not have information about the treatment histories of the patients.

Our study included samples from patients who underwent routine drug resistance testing. In our setting this is mostly performed in heavily treatment experienced patients who are assessed for third-line antiretroviral treatment. However, integrase strand transfer inhibitors (INSTI) have not been used widely in our setting, until recently, which explains the lower rate of scored drug resistance mutations amongst these *IN* sequences. In contrast, among *PR-RT* sequences there were many mutations detected by both sequencing platforms, which allowed us to assess the impact of discordant results. There were no discordant interpretations of 21 cases with *IN* sequencing, whereas for *PR-RT* of 245 Stanford DB mutations detected by either, 238 were detected by both platforms; a large proportions of scored mutations were mixtures – 45 (18.6%) of the 241 detected by ONT and 55 (22.7%) of the 242 detected by Sanger. Only 2 of these 46 *PR-RT* sequences had differences in drug resistance interpretation with in the one Sanger sequencing indicating high level drug resistance to AZT versus intermediate level drug resistance for the same sample according to ONT; and in another case where Sanger sequencing indicated low level ETR resistance whereas ONT indicated potential low level ETR resistance. Considering the slight changes in interpretation it is unlikely that any of these would have resulted in a different regimen choice and these are therefore not likely clinically significant differences. Although another group investigated ONT sequencing in treatment naïve patients[16] ours is the first to assess its performance in samples from treatment experienced patients. Moreover, our results were generated at a low sequence depth (400 read coverage per sample) which makes multiplex testing on affordable Flongle flow cells feasible and may enable one to test much larger batches than the 24 samples on the same flow-cell using combinations of the 96 available ONT barcodes. Many studies have emphasized the ability of next generation sequencing to detect minor drug resistance variants. However, the clinical benefit of this over and above detection of majority variants may be limited to particular settings such as pre-treatment drug resistance assessment or when using etravirine in treatment-experienced patients[17–19]. Moreover, PCR resampling error and sequencing error compound at low frequency variant thresholds so that the lower the threshold the higher the uncertainty of measurement and the more unclear whether these mutations are of clinical significance[13]. Approaches to improve the accuracy such as using random primer IDs, are cumbersome from a workflow and bioinformatics approach[20,21]. When HIV drug resistance testing is performed for the use-case of determining whether drug resistance explains current regimen failure, in deciding whether to switch to another regimen and to inform about the regimen choice, based on guidelines and genotypic susceptibility scores (GSS), Sanger sequencing or an equivalent variant threshold currently remains the standard of care[17].

In future, Nano-RECall will be integrated with ONT basecalling to allow “real-time” reporting of data and QC metrics collected for each sample until reaching a pre-defined read-depth. This would further allow one to optimize coverage and limit run-time, contributing to faster sample-to-result turnaround times.

## Conclusions

Our study has shown that ONT sequencing at relatively low (400 fold) coverage with the Nano-RECall bioinformatics pipeline, which has been designed cognizant of ONT homopolymer read error and which corrects for the resulting alignment errors, can achieve a Sanger equivalent drug resistance interpretation. As a large proportion of drug resistance mutations occur as codon mixtures, this pipeline has benefit over pipelines that provide a majority consensus only. As ONT requires a much lower capital layout than competing platforms and have low sequencing costs when using Flongle flow cells, this may provide an attractive alternative for resource limited settings, that are currently without access to HIV drug resistance sequencing.

The Nano-RECall pipeline is freely available as a downloadable application on a Windows computer at https://recall.hcvdb.ubc.ca/index.html or https://github.com/woodsc/nano_recall and provides accurate Sanger-equivalent consensus sequences and HIV drug resistance interpretation. This combined with a simple workflow and the ability to multiplex multiple samples on Flongle flow-cells, requiring low read coverage, would contribute to making sequencing feasible for resource limited settings.

## Competing interests

No authors had any competing interest.

## Authors’ contributions

RH and CW designed the Nano-RECall pipeline and provided output analyses. KD performed the ONT testing with CG’s help. TN assisted with obtaining the Sanger sequences and amplicons for parallel testing. MC oversaw the Sanger Sequencing. GvZ designed the study. UP provided advice during the study conception and reviewed the manuscript. GvZ wrote the draft manuscript. All authors approved the final manuscript.

## Acknowledgements

We would like to acknowledge the National Health Laboratory Service staff for providing the amplicons and performing the drug resistance tests that were Sanger Sequencing based.

## Funding

The study was funded by Stellenbosch University Division of Virology divisional funds. GvZ received funding from the U.S. National Institutes of Health and South African Medical Research Council through its U.S.-SA Program for Collaborative Biomedical Research (Grant no U01CA200441)

The authors report no conflict of interest.

## Supplement

Additional file 1: Phylogenetic trees of sequences.

## Notes

### Competing Interest Statement

The authors have declared no competing interest.

