## Supplementary figures and images for "An assessment of Nano-RECall: Interpretation of Oxford Nanopore sequence data for HIV-1 drug resistance testing"

### Supplementary figure 1

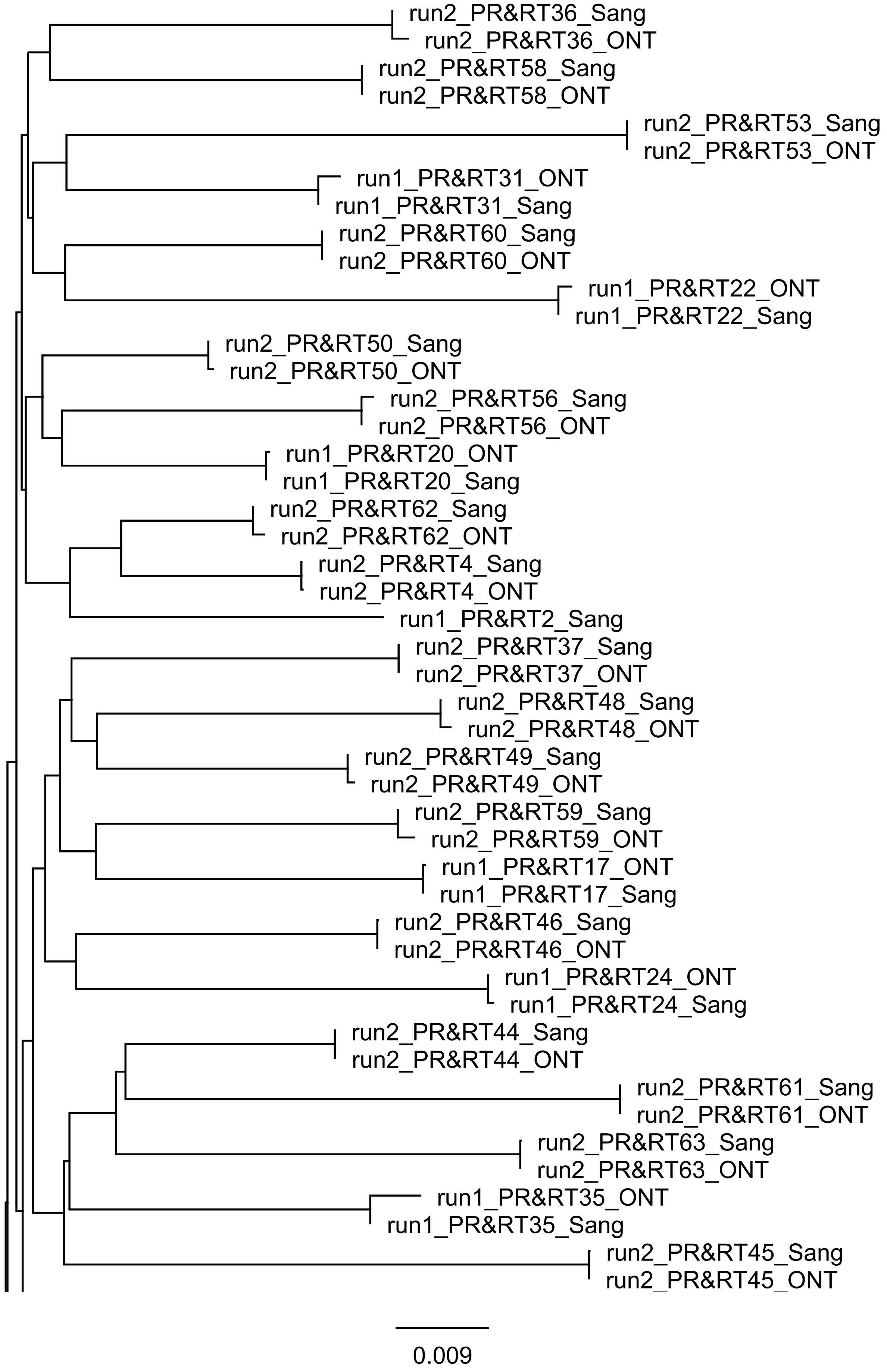

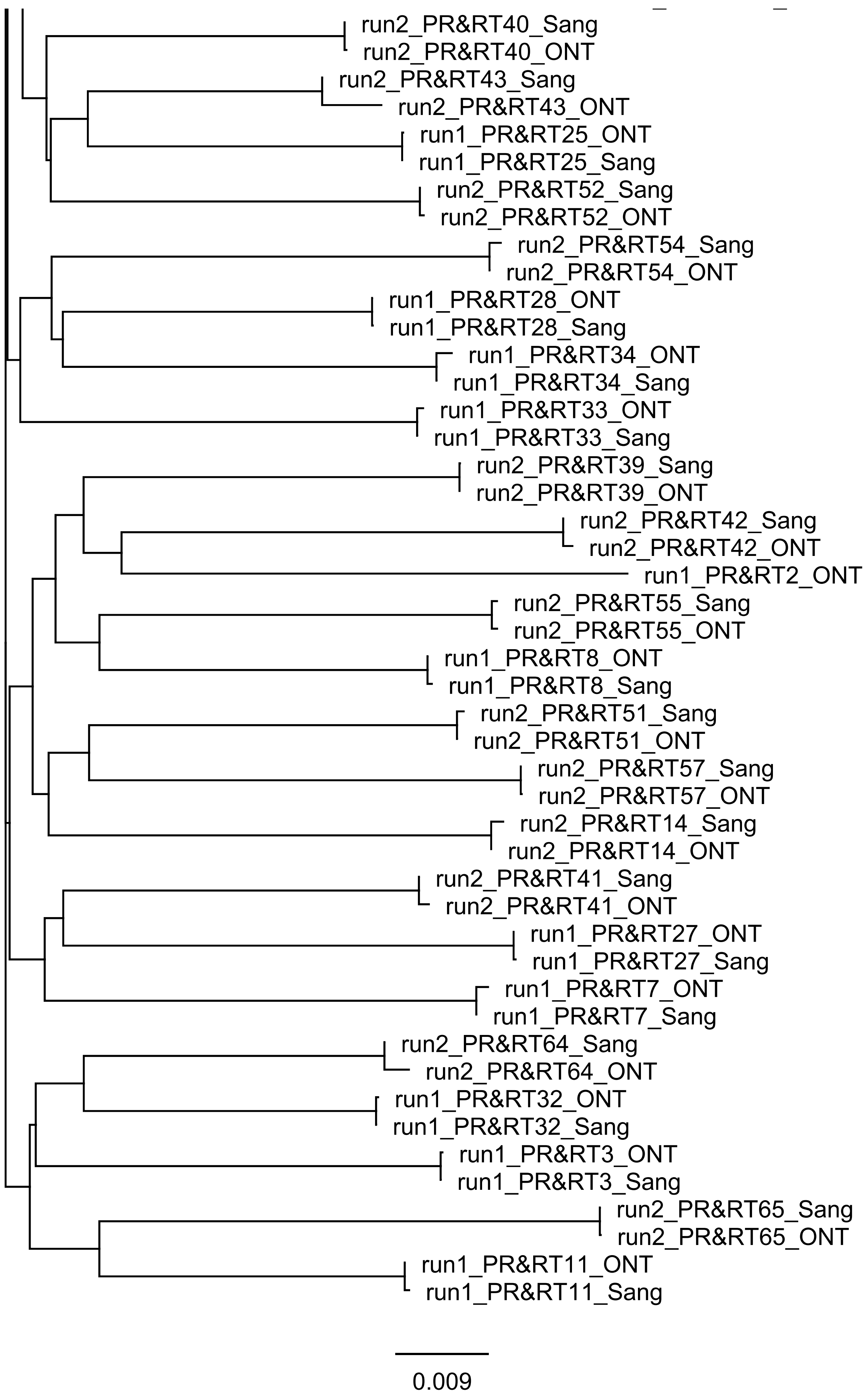

### Supplementary figure 2

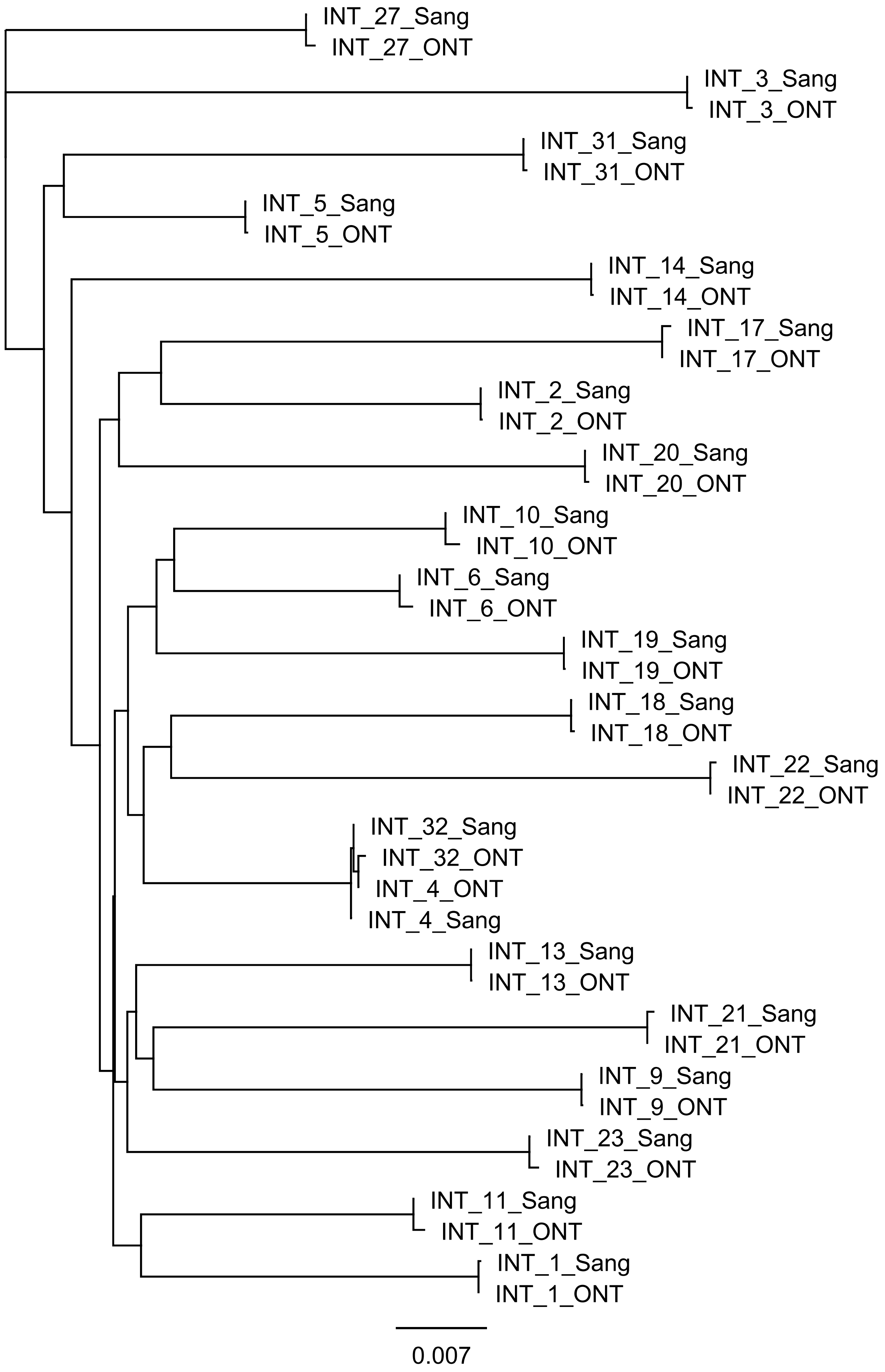
